# Modulating Fis and IHF binding specificity, crosstalk and regulatory logic through the engineering of complex promoters

**DOI:** 10.1101/614396

**Authors:** Lummy Maria Oliveira Monteiro, Ananda Sanches-Medeiros, Cauã Antunes Westmann, Rafael Silva-Rocha

**Author notes:** Correspondence to: Rafael Silva-Rocha, Faculdade de Medicina de Ribeirão Preto, Universidade de São Paulo, Av. Bandeirantes, 3.900. CEP: 14049-900., Ribeirão Preto, São Paulo, Brazil.

## Abstract

Bacterial promoters are usually formed by multiple *cis*-regulatory elements recognized by a plethora of transcriptional factors (TFs). From those, global regulators are key elements since these TFs are responsible for the regulation of hundreds of genes in the bacterial genome. For instance, Fis and IHF are two global regulators which play a major role in gene expression control in *Escherichia coli* and usually multiple *cis*-regulatory elements for these proteins co-occur at target promoters. Here, we investigated the relationship between the architecture of the *cis*-regulatory elements for Fis and IHF in *E. coli*. For this, we constructed 42 synthetic promoter variants harboring consensus *cis*-elements for Fis and IHF at different distances from a core −35/−10 region and in different numbers and combinations. We first demonstrated that although Fis preferentially recognizes its consensus *cis*-element, it can also recognize, to some extent, the consensus binding site for IHF, and the same was true for IHF, which was also able of recognizing Fis binding sites. However, changing the arrangement of the *cis*-elements (i.e., the position or the number of sites) can completely abolish unspecific binding of both TFs. More remarkably, we demonstrate that combining *cis*-elements for both TFs could result in Fis and IHF repressed or activated promoters depending on the final architecture of the promoters in an unpredictable way. Taken together, the data presented here demonstrate how small changes in the architecture of bacterial promoters could result in drastic changes in the final regulatory logic of the system, with important implications for the understanding of natural complex promoters in bacteria and their engineering for novel applications.

**Importance:** The understanding of the regulatory complex in bacteria is a key issue in modern microbiology. Here, we constructed synthetic bacterial promoters in order to investigate how binding of transcriptional factors to multiple target sites at the promoters can influence gene expression. Our results demonstrate in a systematic way that the arrangement and number of these *cis*-regulatory elements are crucial for the final expression dynamics of the target promoters. In particular, we show that TF binding specificity or promiscuity can be modulated using different promoter architectures based on consensus *cis*-regulatory elements, and that transcriptional repression and activation can also be affected by promoter architecture. These results are relevant both for the understanding of natural systems and for the construction of synthetic circuits for biotechnological applications.

## Introduction

Bacteria have evolved complex gene regulatory networks to coordinate the level of expression of each gene in response to changing environmental conditions. In this sense, a typical bacteria such as *Escherichia coli* uses around 300 different transcriptional factors (TFs) to control the expression of its more than 5000 genes, and gene regulation in bacteria has been extensively investigated in the last 6 decades (1). Among the known TFs from *E. coli*, global regulators are those able to control the highest percentage of transcriptional units in response to major physiological or environmental signals, such as the metabolic state of the cell, the availability of carbon sources, the presence of oxygen (2, 3), while local regulators are responsible for gene regulation in response to specific signals (such as sugars, metals) (3, 4). Most TFs control gene expression through their interaction with specific DNA sequences located near the promoter region, the *cis*-regulatory element or TF binding site (4, 5). Over the decades, many *cis*-regulatory elements for many TFs from *E. coli* have been experimentally characterized, mapped and compiled in databases such as RegulonDB and EcoCyc (6, 7). Analysis of these datasets demonstrates that TFs usually act in a combinatorial way to control gene expression, where multiples *cis*-regulatory elements for different TFs are located in the upstream region of the target genes (6, 8, 9). Therefore, the arrangement of *cis*-regulatory elements at the target promoters is crucial to determine which TFs will be able to control the target gene and how these regulators will interact among each other once bound to the DNA (3, 10).

Several previous works have explored the relationship between the architecture of *cis*-regulatory elements and the final logic of the target promoters, and initial attempts on this sense have focused on the mutation of *cis*-regulatory elements from natural promoters to investigate how these elements specify the promoter expression dynamics (11–14). More recently, Synthetic Biology approaches have been used to construct artificial promoters through the combination of several *cis*-regulatory elements, and these have been characterized to decipher their architecture/dynamics relationship (15–18). Yet, while most Synthetic Biology approaches have focused on *cis*-elements for local regulators (which not commonly regulate gene expression in a combinatorial way), we recently investigated this combinatorial regulation problem with global regulators (8, 19, 20). This is important since global regulators (such as IHF, Fis, CRP among others) have numerous binding sites along the genome of *E. coli* and very frequently co-occur at target promoters (8). In this sense, Fis and IHF are two global regulators with critical role in coordinating gene expression in *E. coli* as well as in mediating DNA condensation in the cell (3, 4, 21, 22). Fis, a very abundant nucleoid associated protein (NAP) is related to gene expression regulation in fast-growing cells, while IHF is a NAP related to changes in gene expression in cells in the transition from exponential to stationary phase (21, 23, 24). Moreover, IHF binds to AT rich DNA motifs with well-defined sequence preferences (25, 26), while Fis has also preference for AT reach regions with a more degenerated sequence preference (27, 28). Additionally, cross-regulation between Fis and IHF have been demonstrated for several systems (3, 21, 29), and how specific vs. promiscuous DNA recognition can be achieved for these two global regulators is not fully understood.

We previously explored how complex synthetic promoter harboring *cis*-regulatory elements for CRP and IHF can generate diverse regulatory logics depending on the final architecture of synthetic promoters, demonstrating that it is not possible to predict the regulatory logic of complex promoters from the known dynamics of their simple versions (19). Here, we further explore this approach to investigate the relationship of *cis*-regulatory elements for Fis and IHF. Using consensus TF binding sites for these two TFs at different promoter positions and in different numbers, we first demonstrated that while some promiscuous interactions occurs between the TFs and the binding sites, some specific *cis*-regulatory architectures can completely abolish unspecific interactions. Additionally, and unexpectedly, complex promoters constructed by the combination of *cis*-elements for Fis and IHF can generate many completely different outputs, such as Fis-repressed promoters, IHF-repressed promoters or systems where Fis and IHF act as activators. As these changes in promoter logic results from changes in promoter architecture only (and not on the affinity of the TF to each individual *cis*-elements), the data presented here reinforce that notion that complex bacterial promoters can display emergent properties, where their final behavior cannot be defined from the characterization of the individual component. Taken together, the data presented here provide insightful evidence for the understand of natural complex promoters controlled by global regulators, as well as has implication for the forward engineering of synthetic promoters.

## Results and discussion

### Creating complex promoters for Fis and IHF

In order to investigate the effect of promoter architecture in the regulation by Fis and IHF, we constructed a number of combinatorial promoters where consensus DNA sequences for Fis and IHF binding are upstream of a weak core promoter (−35/−10 region) at specific positions (1 to 4), which are centered at the −61, −81, −101 and −121 regions related to the transcriptional start site (TSS) (Fig. 1). In this sense, double stranded DNA sequences were generated for each position and a control sequence (Neg), to which no TF can bind, was used a control (20). After selecting consensus sequences for each TF, the complex promoters were assembled by DNA ligation and cloned into a mid-copy number vector harboring two reporter genes for fluorescent proteins (mCherry and GFPlva, Fig. 1). The resulting reporter plasmids (with each promoter controlling GFPlva expression) were used to transform competent wild-type strains or *ihf* or *fis E. coli* mutants (30). Using this approach, we could assay promoter activity measuring GFP production in all strains in a plate reader during growth in minimal media for 8 hours. Therefore, we created a library of 42 complex promoters containing different number of copies of *cis*-regulatory elements for single TFs or mixed binding sites for both Fis and IHF (Table 1). In the next sections, we presented the results of the promoter analysis per categories to uncover the *cis*-regulatory logic for each variant.

**Table 1.**
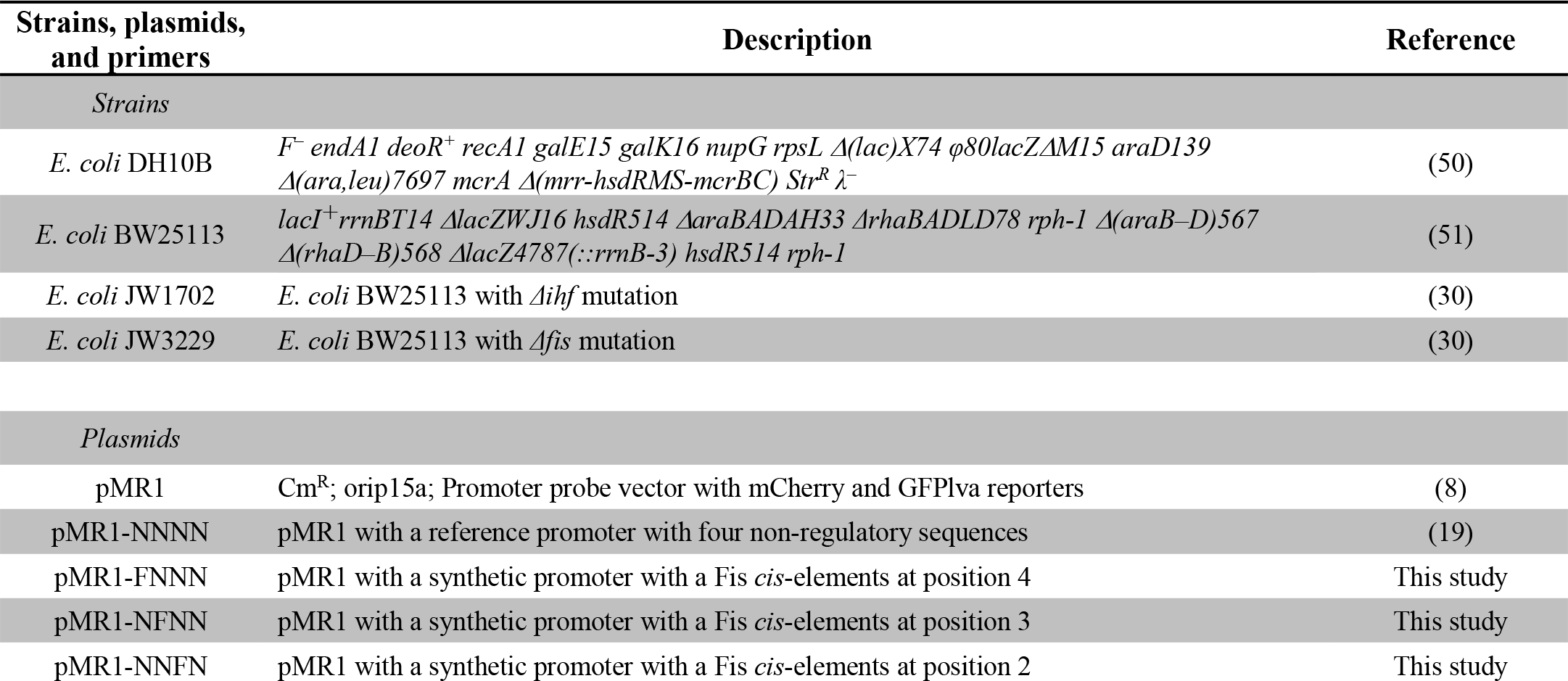
Strains, plasmids and primers used in this study.

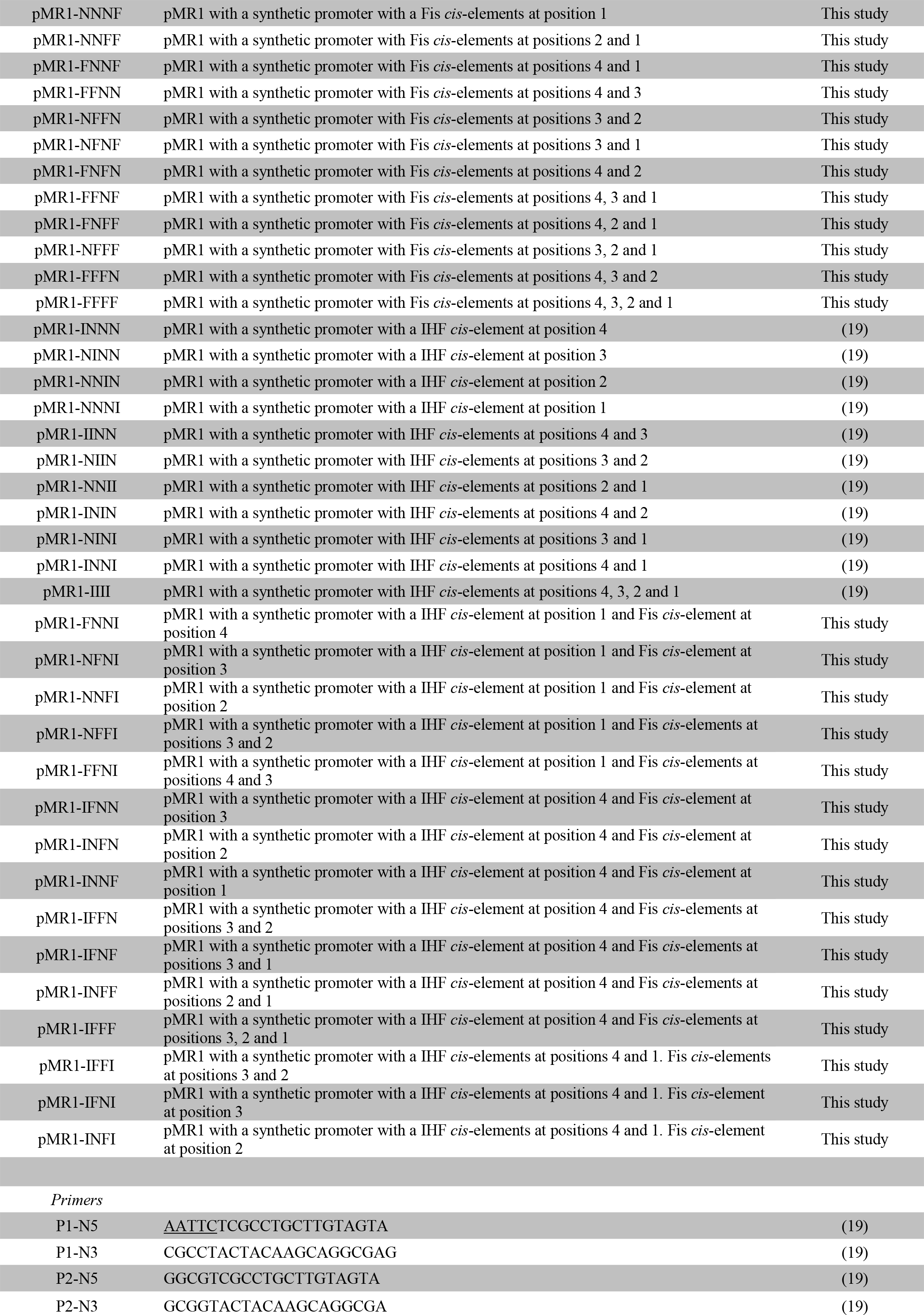

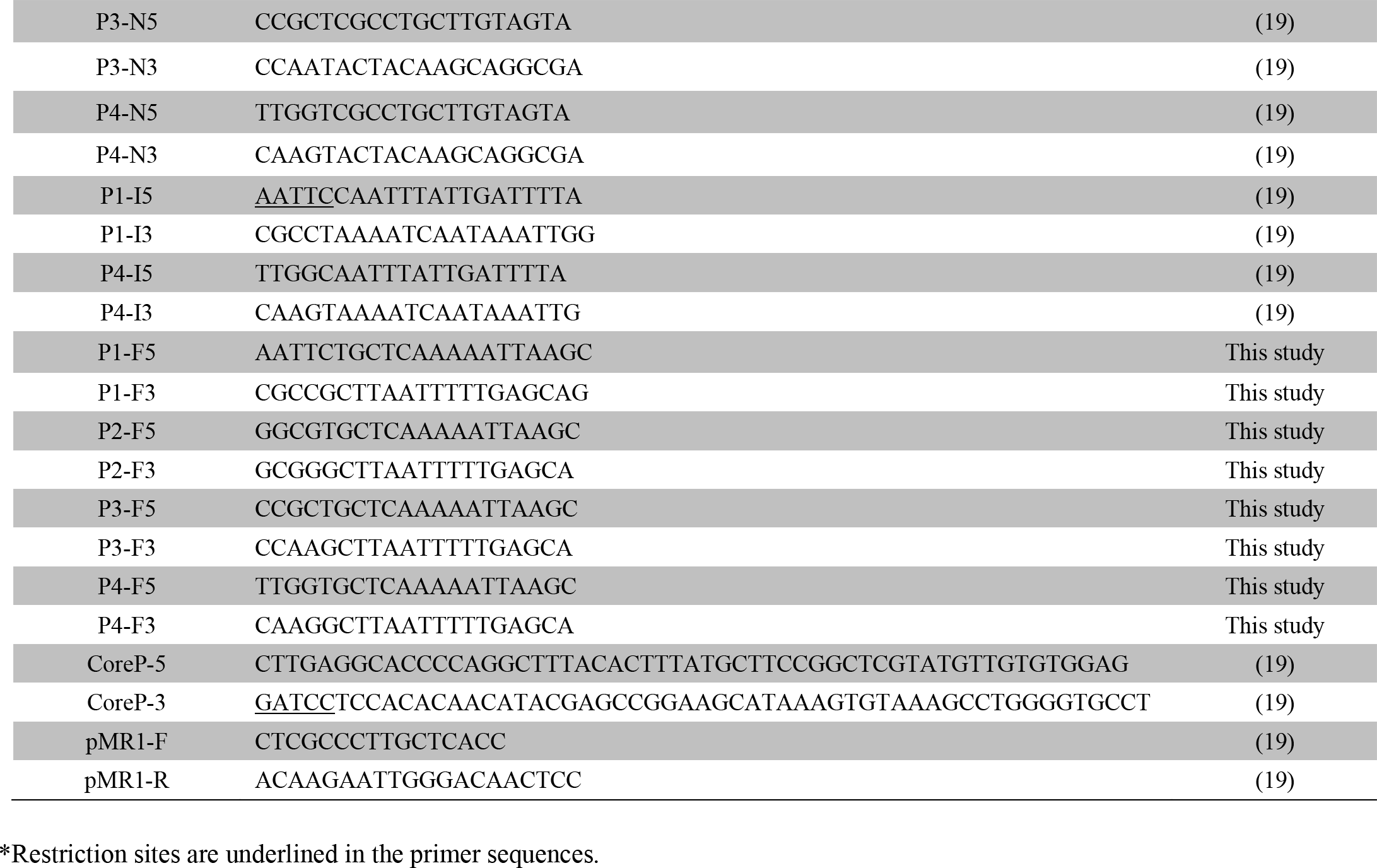

**Figure 1.**
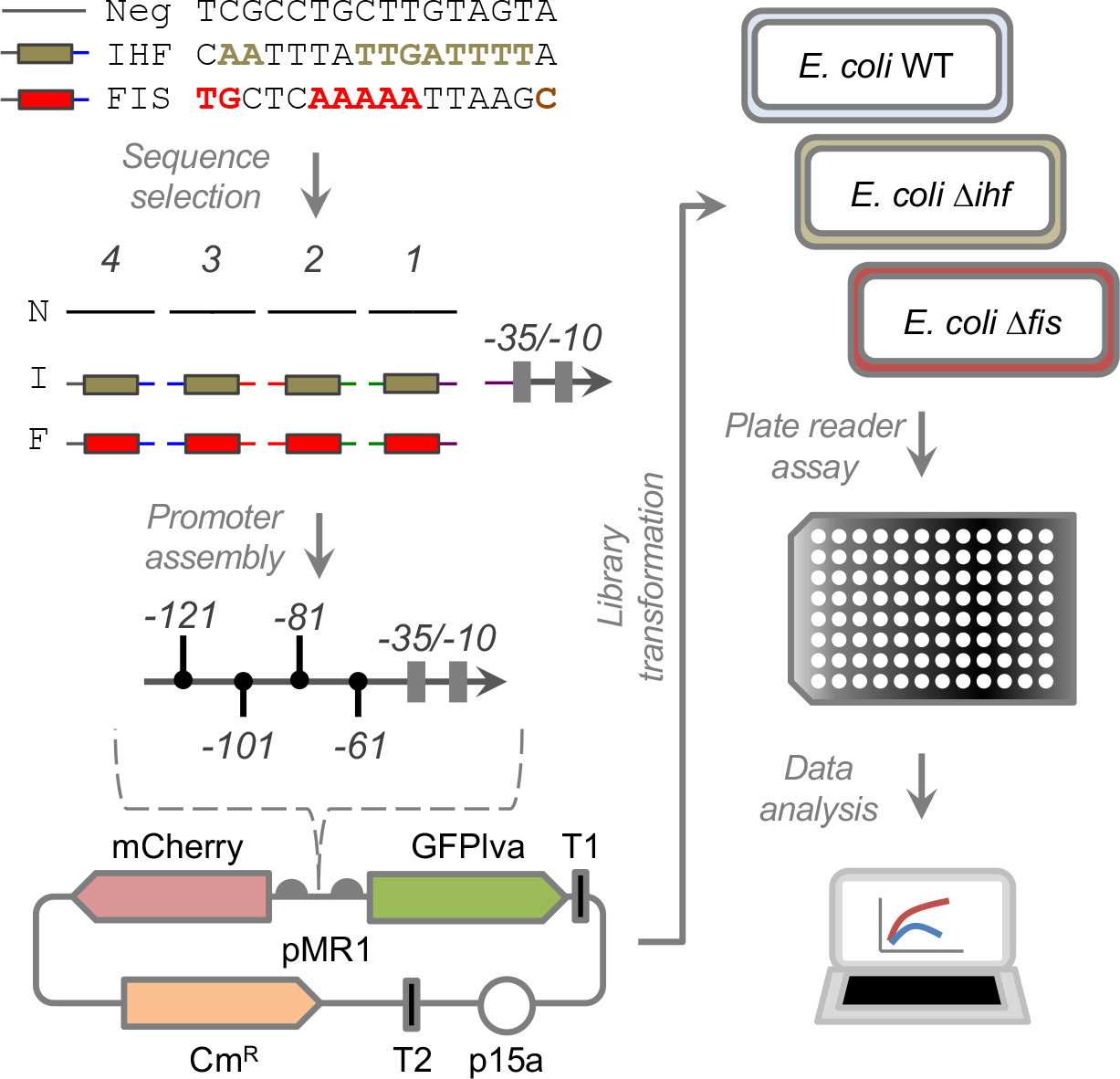
Strategy to construct synthetic complex promoters. DNA sequences harboring the consensus sequence for IHF or Fis binding were selected, along with a control sequence each cannot be recognized by any TF. Double stranded DNA fragments were produced with cohesive ends specific for each promoter positions (numbered from 1 to 4) and assembled together with a weak core promoter harboring the −35/−10 boxes for RNAP recognition (REF). The fragments were cloned into a promoter probe vector (pMR1) harboring resistance to chloramphenicol (Cm^R^), a medium-copy number origin of replication (p15a) and two reporter genes (mCherry and GFPlva). The libraries were introduced into wild type and mutant strains of *E. coli* from the KEIO collection (30). The resultant strains were analyzed at the population level in a plate reader and the data processed using script in R.

### Changing Fis binding sites architecture modulates Fis and IHF binding specificity

We started by analyzing the effect of binding site copy number and arrangement of *cis*-regulatory elements for Fis. For this, we constructed several promoter variants and assayed their dynamics in wild-type, *Δfis* and *Δihf* strains of *E. coli*. As can be seen in Fig. 2, most promoters constructed produced very low activity over the growth of wild type *E. coli*. However, when these promoters were assayed in the *E. coli Δfis* strain, four promoter variants harboring one, two or three binding sites for Fis displayed significant increase in activity (promoters shaded in blue and orange in Fig. 2). These results indicated that Fis was acting as a repressor of promoter activity for these four variants. Next, we assayed promoter activity in *E. coli Δihf* strain, in order to see if IHF could also exert some regulatory interaction with the promoters harboring binding sites for Fis. As can been seen in Fig. 2, most promoters displayed similar expression levels as when in the wild type strains of *E. coli*. Yet, a single promoter variant harboring a Fis binding site at position 3 (−101 relative to the TSS) displayed a strong increase in activity relative to the wild type strain, indicating that IHF is acting as a repressor of this promoter variant (promoter shaded in orange in the Fig. 2). All together, these results indicated that while unspecific IHF binding to the Fis *cis*-regulatory element exists, it was restricted to a single promoter variant, suggesting that promiscuous regulatory interaction can be avoided by changing promoter architecture.

**Figure 2.**
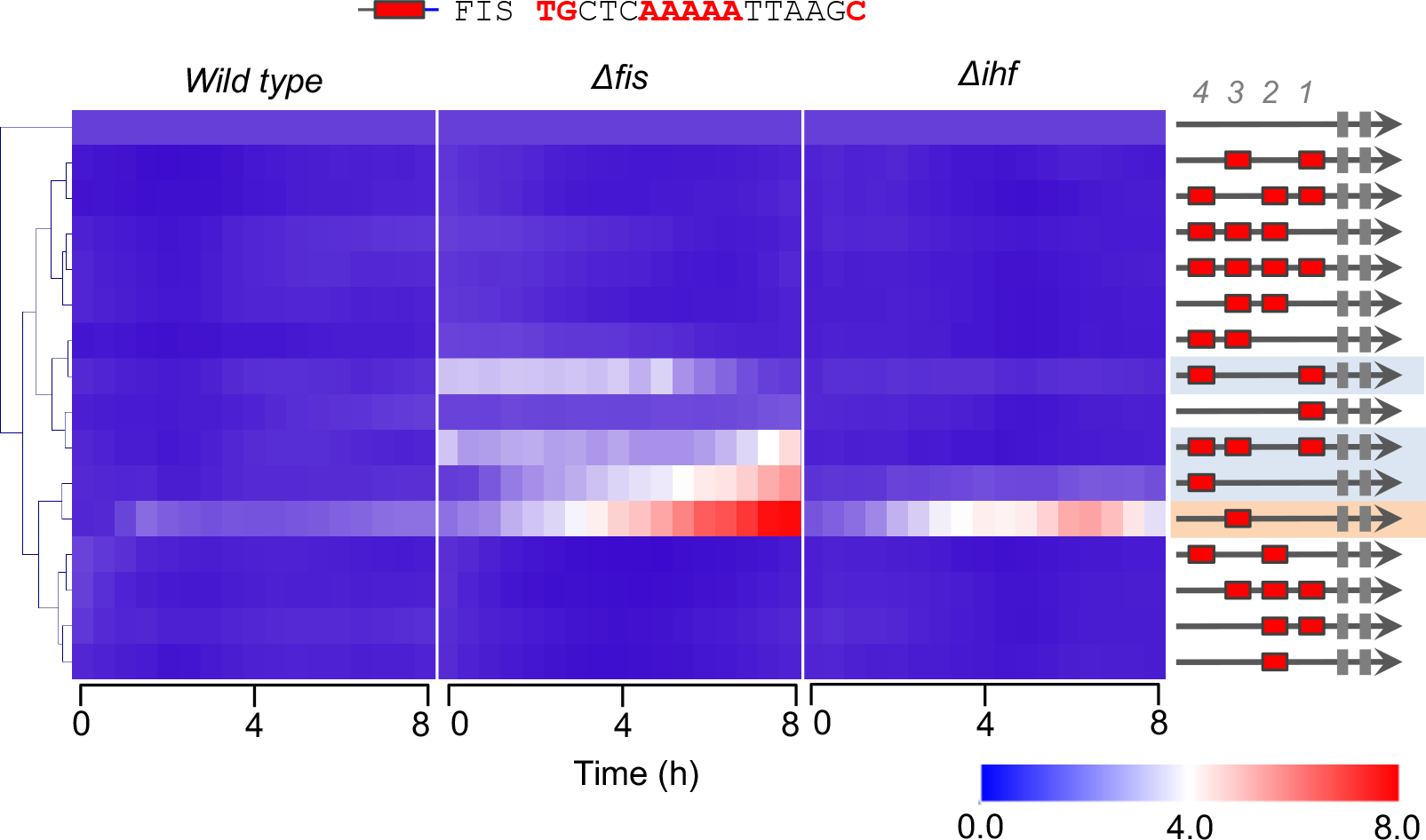
Activity of promoters harboring Fis-binding sites. The architecture of the synthetic promoter is shown on the left. Promoters activities are shown in heatmaps and normalized pee the activity of the reference promoter (i.e., a promoter with four control sequences). Promoter analysis were monitored over 8 hours with measurements every 30 min in three genetic backgrounds of *E. coli* (wild type, *Δfis* and *Δihf*). The results shows the average of three independent experiments. Promoters are clustered using the Euclidian distance using MeV software.

### IHF binding sites can be recognized by Fis regulator in an architecture-dependent manner

We next investigated the effect of Fis and IHF in the regulation of promoters harboring multiple cis-regulatory elements for IHF. As can be seen in Fig. 3, most promoters assayed displayed low activity in wild type strain of *E. coli* and higher activity in the mutant strain lacking *ihf*, in agreement with previous data on complex IHF promoters (19). However, when these promoters were assayed in *E. coli Δfis* strain, we observed that three promoter architectures also displayed higher activity in the mutant (promoters shaded in blue in the figure), indicating that Fis was also able to repress these promoter variants. Yet, it is worth noticing that a promoter variant harboring two *cis*-regulatory elements for IHF at positions 3 and 4 (−101 and −121 relative to the TSS) displayed both a strong repression by IHF but no modulation by Fis (promoter shaded in orange in the figure). Again, these results reinforce the notion that promiscuous or specific TF binding can be modulated by changing the promoter architecture.

**Figure 3.**
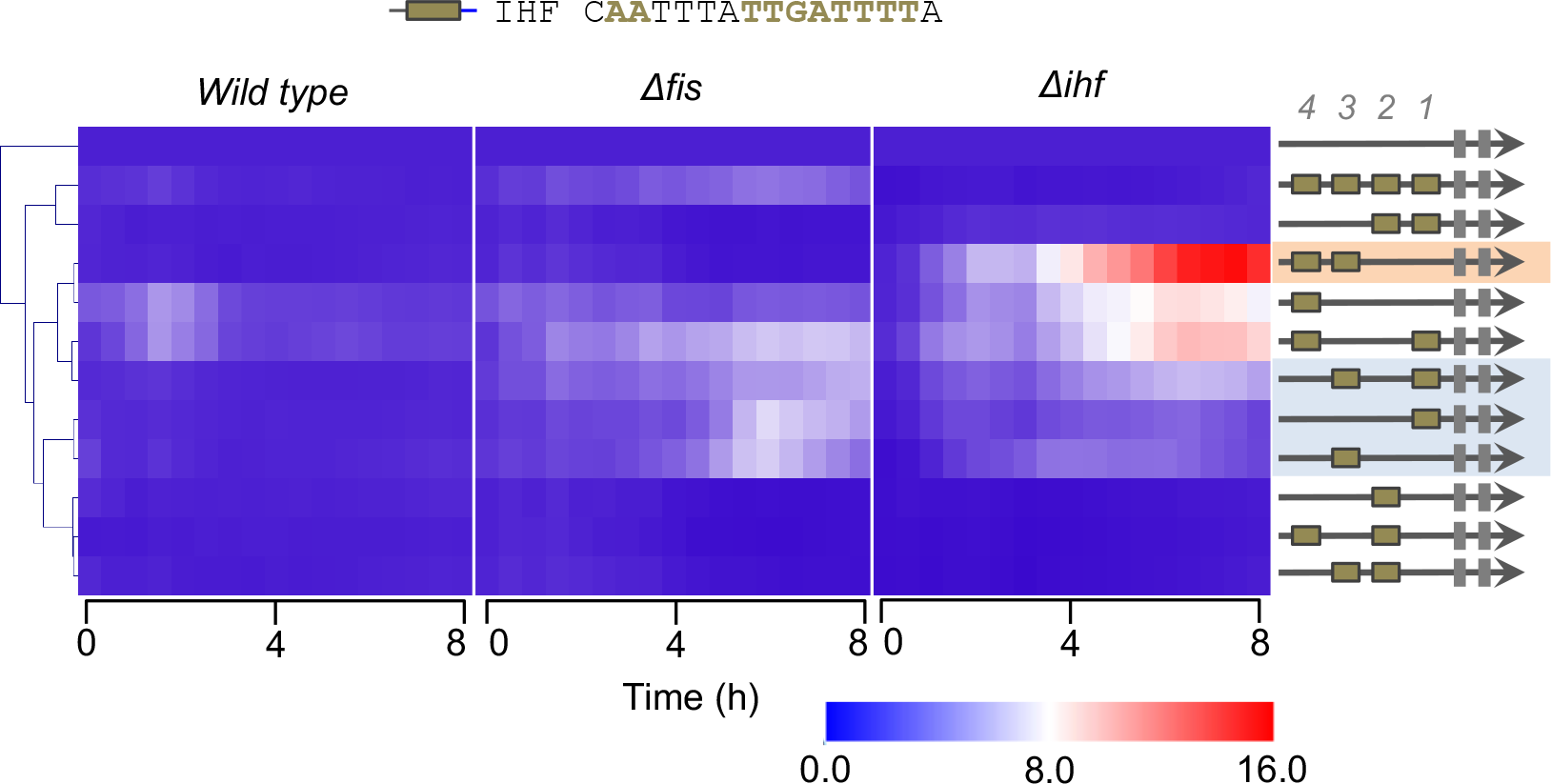
Activity of promoters harboring IHF-binding sites. Experiments were preformed and data presented as in Fig. 2. 11 promoters variants previously described (19) were analyzed in wild type, *Δfis* and *Δihf* mutant strains of *E. coli*.

### Intrinsic DNA architecture modulates promoter activity independently of regulator binding

Once we investigated the regulatory interactions for promoters harboring *cis*-regulatory elements for a single TF (IHF or Fis), we constructed promoters with binding sites for both TFs. In order to systematically investigate the effect of combined TF-binding sites on promoter logic, we first placed an IHF-binding site at position 1 (−61) and varied Fis-binding sites number and positions on these promoters. As shown in Fig. 4A, a promoter harboring a single IHF-binding site at position 1 presented no activity in wild type strain of *E. coli* but they increase activity in the *fis* and *ihf* mutant strains. However, adding Fis-binding sites at position 2 or 3 resulted in promoters with reduced activity even in the *ihf* mutant strain, and this repression was not alleviated in the *fis* mutant strain (promoters shaded in blue in Fig. 4A). This indicate that the repression of the promoters harboring additional Fis-binding sites should be due to the intrinsic DNA geometry of the resulting promoter, as reported previously for other systems (31, 32). When the single IHF-binding site was fixed at position 4 (−121), the resulting promoter displayed strong activity in wild type and *Δihf* strains, as well as increased activity in *Δfis* strain (Fig. 4B). However, when a single Fis *cis*-regulatory element was added at position 1 (−61), the resulting promotor displayed increased activity in *E. coli Δfis* strain, while it presented no activity in wild type and *Δihf* strain. In this sense, the new promoter was now repressed only by the Fis regulator. Finally, the addition of a single or multiple Fis-binding sites at different positions completely blocked promoter activity, and this was not relieved either in *Δfis* or *Δihf* strains. Taken together, these results also suggest that DNA geometry can interfere with promoter activity in a TF-independent way, as most combinatorial promoters shown in Fig. 4B displayed no detectable activity.

**Figure 4.**
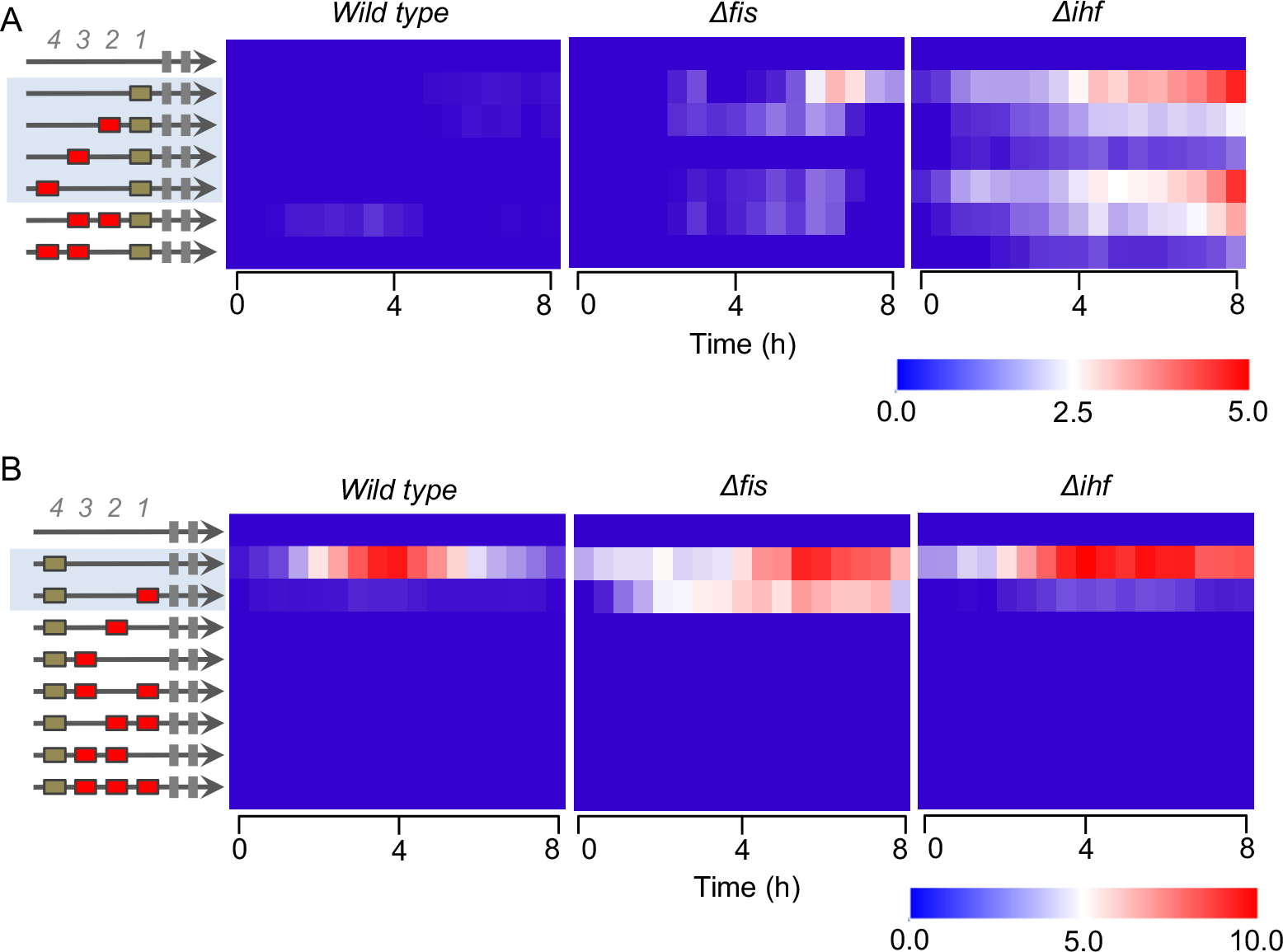
Activity of promoters with combined IHF- and Fis-binding sites. All experiments were performed and analyzed as in Fig. 2. **A)** Characterization of promoters with a single IHF-binding site fixed a position 1 (−61) and varying Fis-binding sites. **B)** Characterization of promoters with a single IHF-binding site fixed a position 4 (−121) and varying Fis-binding sites.

### Combination of Fis and IHF binding sites generates strong Fis and IHF activated promoters

In all promoters presented until this point, while the combination of different *cis*-regulatory was able to determine the regulatory logic displayed by IHF and Fis, the two TFs were acting as repressors of promoter activity (Figs. 2–4). Yet, this behavior shifted when we constructed promoters version harboring IHF-binding sites at positions 1 and 4 and varying sites for Fis (Fig. 5). As shown in this figure, when a single Fis-binding site was added at position 2 (−81), the resulting promoter displayed a strong activity in the wild type strain of *E. coli*, when compared to the version lacking this element (promoters in the blue shaded region at Fig. 5). Furthermore, when these promoters were assayed in the mutant strains lacking Fis or IHF, we could observe a strong reduction in their activity, indicating that both TFs were acting as activators of the combinatorial promoters. The same partner was also observed for a promoter harboring the two IHF-binding sites (at position 1 and 4) and two Fis-binding sites (positions 2 and 3). These unexpected results highlight rise of emergent properties in complex promoters for global regulators (19), as increasing the number of *cis*-regulatory elements can drastically shift the final regulatory logic of the system.

**Figure 5.**
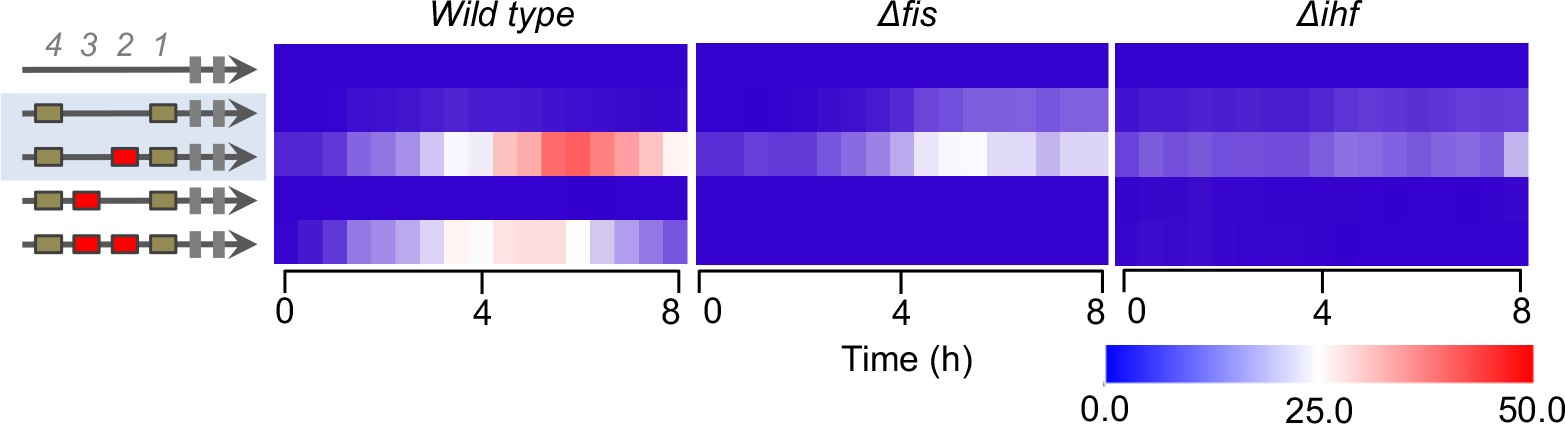
Analysis of promoters with two fixed IHF-binding sites. For this analysis, IHF *cis*-regulatory elements were placed at positions 1 and 4 and additional Fis-binding sites were introduced into the promoters. All promoters were assayed in wild type, *Δfis* and *Δihf* mutant strains of *E. coli*.

### Conclusions

Bacteria are naturally endowed with complex promoters harboring multiple binding sites for several TFs. While several works based on mathematical modelling have argued that combinatorial regulation can be predicted from the characterization of individual promoter elements (33–36), we are providing here and previously (19) growing evidence that small changes in the architecture of *cis*-regulatory elements can drastically change the final response of the system (37). The unpredictable behaviors observed in these works might also depict a deeper evolutionary trend in gene regulation that has selected molecular systems/mechanisms capable of promoting both evolvability and robustness of gene expression levels through nonlinear gene regulation (38). Thus, understanding the way the architecture of *cis*-regulatory elements determine gene expression behavior is pivotal not only to understand natural bacterial systems, but also to provide novel conceptual frameworks for the construction of synthetic promoters for biotechnological applications. Moreover, we were able to grasp, even in a narrow-scale subset of combinatorial diversities, biologically relevant effects of transcriptional crosstalk, a phenomenon that has been widely explored in eukaryotes, but overlooked in prokaryotes due to their significantly longer TF binding motifs with higher information-content (39, 40) – suggesting a predicted lower probability of crosstalk (41, 42). Theoretical studies (41–48) have suggested important biological and evolutionary roles and tradeoffs for transcriptional crosstalk such as imposing intrinsic costs to cellular systems (41, 42, 44, 46) – as gene expression might be improperly activated or repressed by non-cognate TFs at specific environmental contexts –, while also generating evolutionary flexibility that might be beneficial during the evolution of novel regulators through gene duplication (44). Theoretical models have also been recently developed, suggesting the modulation of transcriptional crosstalk by different regulatory logic architectures (41, 44), a phenomenon that could be experimentally supported in the present work. Future studies, validating alternative combinatorial subsets of promoters should greatly improve the comprehension of the rules underlying transcriptional crosstalk modulation, with direct impact on the understanding and engineering of biological systems. Fig. 6 provides a visual summary of some of the finds reported here under a Boolean logic perspective. As shown in Fig. 6A, changing a perfect Fis binding in 20 bp (from position −121 to −101) can turn a specific Fis-repressed promoter into a system repressed by both Fis and IHF. Using a more formal logic gate definition (49), this modification can turn a promoter with a NOT logic into one with a NOR logic. On the other hand, a promoter harboring two IHF-binding sites at positions −121 and −101 displayed specific IHF-repression, while changing the second binding site to position −61 resulted in a promoter repressed by both IHF and Fis (Fig. 6B). In terms of promoter logic, this change in *cis*-element architecture also turn a promoter with NOT logic into one with a NOR logic. When a single IHF-binding site was presented at position −121, the final promoter was only repressed by IHF (Fig. 6C). Yet, introducing an additional Fis-binding site at position −61 of this promoter turned it into a system exclusively repressed by Fis. This change maintained the NOT logic of the promoter but changed the TF able to repressed it activity. Finally, and more remarkably, while a promoter with two IHF-binding sites (at positions −121 and −61) were repressed by both Fis and IHF, adding a third binding site for Fis at position −81 resulted in a promoter strongly activated by both TFs (Fig. 6D). Therefore, this single change *cis*-element architecture turned a promoter with NOR logic into a fully OR promoter responsive to the same TFs. This remarkable regulatory versatility and unpredictability unveiled by synthetic combinatorial promoters evidences that we are still starting to understand how complex gene regulation in bacteria can be. While the work presented here cover two of the main global regulators of *E. coli*, further works are still necessary to uncover the hidden complexity of combinatorial gene regulation in this bacterium.

**Figure 6.**
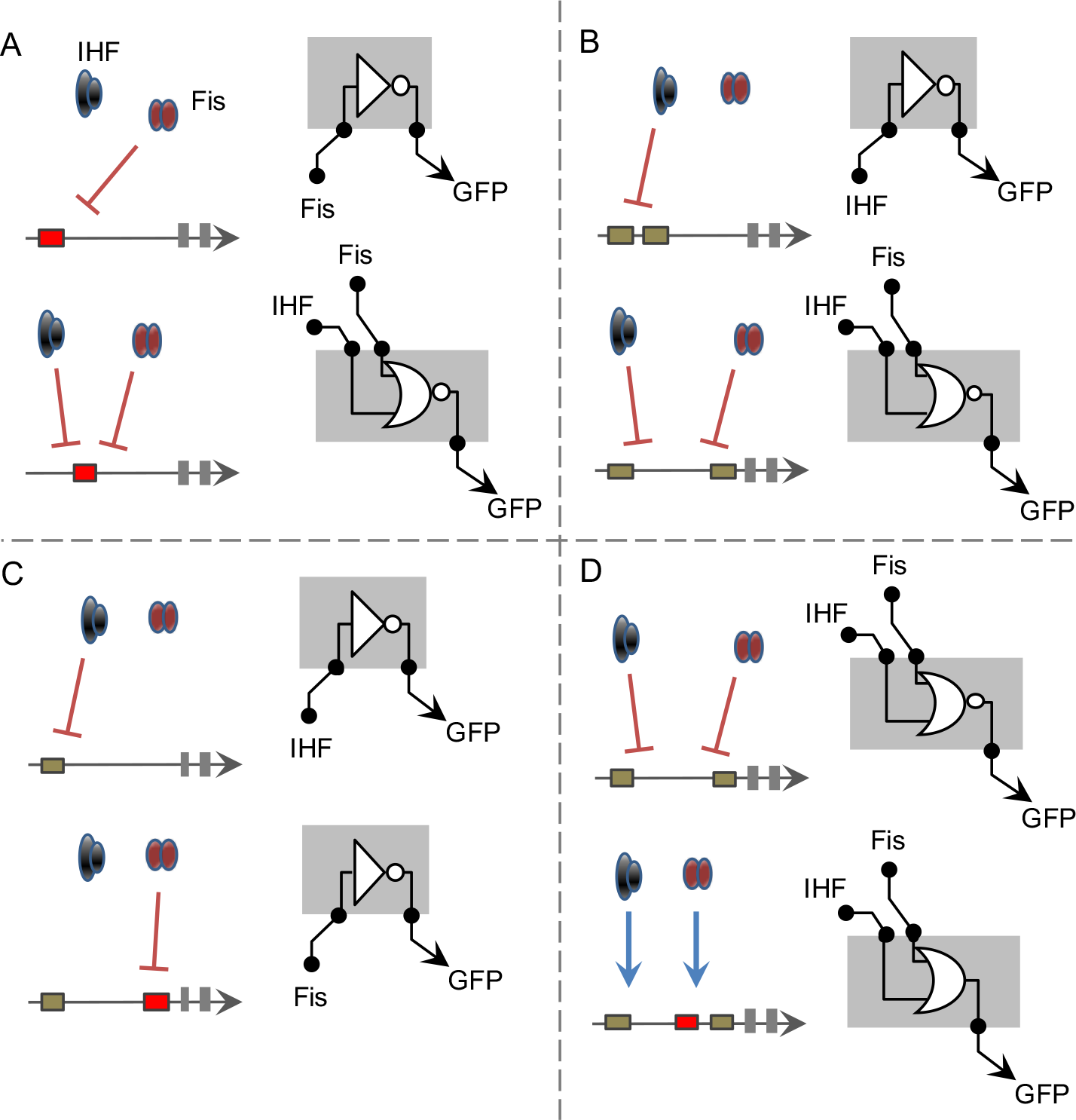
Summary of most significant changes in promoter architectures leading to changing in promoter logic. The figures are represented using logic gate representation for gene regulation even (49). **A)** the change of a single Fis-binding site from position −121 to −101 turns a NOT gate for Fis into a NOR gate for Fis and IHF. **B)** Two IHF-binding sites (at positions −121 and −101) worked as a NOT gate for IHF, while changing one site to position −61generates a NOR gate for both IHF and Fis. **C)** While a single IHF-binding site at position −121 results into a NOT gate for IHF, adding a Fis-binding site to the position −61 creates a NOT gate exclusively dependent on Fis. **D)** Finally, adding a Fis-binding site (position −81) to the NOR gate promoter presented in B drastically changes its logic to a OR gate, were the two TFs acts as activators.

## Material and Methods

### Plasmids, bacterial strains, and growth conditions

*E. coli* DH10B was used for cloning procedures, while *E. coli* BW25113 was used as wild type strain (WT), *E. coli* JW1702-1 was used as mutant for the IHF transcription factor and *E. coli* JW3229 was used as mutant for Fis transcription factor. All strains were obtained from the Keio collection (30). For the procedures and analyses *E. coli* strains were grown in M9 minimal media (6.4 g L^−1^ Na_2_HPO_4_•7H_2_O, 1.5 g L^−1^ KH_2_PO_4_, 0.25 g L^−1^ NaCl, 0.5 g L^−1^ NH_4_Cl) supplemented with chloramphenicol at 34 μg mL^−1^, 2mM MgSO_4_, 0.1mM casamino acids and 1% glycerol as sole carbon source at 37°C. Plasmids, bacterial strains, and primers used in this work are listed in Table 1.

### Design of synthetic promoters scaffold and ligation reactions

The construction of synthetic promoters was performed by ligation reaction of 5’ end phosphorylated oligonucleotides (19) acquired from Sigma Aldrich (Table 1). The design of all single strand was projected to be located at −61, −81, −101 or −121 bp upstream of the core promoter (Fig 1A) and to carry 16 pb sequence containing the Fis binding site (F), IHF binding site (I) or a Neutral motif (N), which is a sequence that any transcription factor is able to bind (Fig. 1B). These locations were identified as position 1, 2, 3, and 4, respectively (Fig. 1C). In addition to the 16 pb oligonucleotides, all single strand was designed to contain three base pairs overhang for its corrected insertion on the promoter (Fig.1A). Additionally, a core promoter based on the lac promoter, which is a weak promoter and therefore requires activation. To perform the assembly of the synthetic promoters, the 5′ and 3′ strand corresponding for each position where mixed at equimolar concentrations and annealed by heating at 95 °C for 5 minutes, followed by gradual cooling to room temperature for 5 minutes and finally it was maintained at 0 °C for 5 minutes. The external overhangs of the *cis*-element at the position four and the core promoter were designed to carry EcoRI and BamHI digested sites, in this way, it is allowed to ligate to a previously digested EcoRI/BamHI pMR1 plasmid. All five fragments (four *cis*-elements positions plus core promoter) were mixed in equimolarity in a pool with the final concentration of 5’ phosphate termini fixed in 15 μM. For the ligase reaction, 1μL of pool of fragments was added to 50 ng EcoRI/BamHI pMR1 digested plasmid in presence of ligase buffer and ligase enzyme to final volume of 10 μL. The ligation was performed for one hour at 16 °C and after that, ligase reaction was inactivated for 15min at 65°C. 2μL of the ligation was used to electroporated 50 μL of *E. coli* DH10B competent cell. After one hour regenerating in 1mL LB media, total volume was plated in LB solid dishes supplemented with chloramphenicol at 34 μg mL-1. Clones were confirmed by colony PCR with primers pMR1-F and pMR1-R (Table 1) using pMR1 empty plasmid PCR reaction as further length reference on electrophorese agarose gel. Clones with potential correct length were submitted to Sanger DNA sequencing for confirmation of correct promoter assembly.

### Promoter activity analysis and data processing

The promoter activity was measured for all 42 promoters analyzed for different genetic backgrounds and conditions. For each experiment, the plasmid containing the interest promoter was used to transform *E. coli* wild type, *E. coli* Δ*ihf* mutant or *E. coli* Δ*fis* mutant as indicated. Freshly plated single colonies were growing for 16 hours in M9 media and then 10μL of this culture was assayed in 96 wells microplates in biological triplicate with 190 μL of M9 media. Cell growth and GFP fluorescence were quantified using Victor X3 plate reader (PerkinElmer) that were measured during 8 hours in intervals of 30 minutes. Promoters activities were calculated as arbitrary units dividing the GFP fluorescence levels by the optical density at 600nm (reported as GFP/OD600) after background correction. Technical triplicates and biological triplicates were performed in all experiments. Row data were processed using *ad hoc* R script (https://www.r-project.org/) and plots were constructed using R or MeV (www.tm4.org/mev.html). For all analyses, the strain under analysis containing pMR1 plasmid was used as threshold background.

## Acknowledgments

This work was supported by the Sao Paulo Research Foundation (FAPESP, award # 2012/22921-8 and 2017/50116-6). LMOM, AS-M and CAW were supported by FAPESP PhD and Master Fellowships (award # 2016/19179-9, 2018/04810-0 and 2016/05472-6). The authors are thanks to lab colleagues for insightful comments on this work.

